# A stomatal model of anatomical tradeoffs between gas exchange and pathogen colonization

**DOI:** 10.1101/871228

**Authors:** Christopher D. Muir

## Abstract

Stomatal pores control both leaf gas exchange and are one route for infection of internal plant tissues by many foliar pathogens, setting up the potential for tradeoffs between photosynthesis and defense. Anatomical shifts to lower stomatal density and/or size may also limit pathogen colonization, but such developmental changes could permanently reduce the gas exchange capacity for the life of the leaf. I developed and analyzed a spatially explicit model of pathogen colonization on the leaf as a function of stomatal size and density, anatomical traits which partially determine maximum rates of gas exchange. The model predicts greater stomatal size or density increases the probability of colonization, but the effect is most pronounced when the fraction of leaf surface covered by stomata is low. I also derived scaling relationships between stomatal size and density that preserves a given probability of colonization. These scaling relationships set up a potential anatomical conflict between limiting pathogen colonization and minimizing the fraction of leaf surface covered by stomata. Although a connection between gas exchange and pathogen defense has been suggested empirically, this is the first mathematical model connecting gas exchange and pathogen defense via stomatal anatomy. A limitation of the model is that it does not include variation in innate immunity and stomatal closure in response to pathogens. Nevertheless, the model makes predictions that can be tested with experiments and may explain variation in stomatal anatomy among plants. The model is generalizable to many types of pathogens, but lacks significant biological realism that may be needed for precise predictions.

## INTRODUCTION

Stomata evolved to regulate gas exchange in and out of the leaf (Hetherington and Woodward, 2003; Berry et al., 2010; Chater et al., 2017), but many foliar pathogens take advantage of these chinks in the leaf cuticular armor to infect prospective hosts (Zeng et al., 2010; McLachlan et al., 2014; Melotto et al., 2017). The stomatal and mesophyll conductance to CO_2_ are two major limits to photosynthesis (Lawson et al., 2018; Flexas et al., 2018) that are partially determined by stomatal anatomy. Since CO_2_ conductance limits photosynthesis (Farquhar and Sharkey, 1982; Jones, 1985) and pathogen infection can reduce fitness (Gilbert, 2002), this sets up a potential tradeoff between increased photosynthesis and defense against pathogens mediated by stomatal anatomy (McKown et al., 2014; Tateda et al., 2019; Dutton et al., 2019; Fetter et al., 2019). For example, plants could increase photosynthetic rate by developing more stomata, but more stomata could result in more pathogen colonization. The optimal stomatal density, size, and arrangement on the leaf will depend on the fitness gains from increased gas exchange and fitness losses imposed by foliar pathogens, both of which depend on the environment. In the next two paragraphs I will review the relationship between stomatal anatomy, gas exchange, and foliar pathogen colonization. Then I will discuss why two anatomical traits, stomatal size and density, might be crucial components of a broader tradeoff between photosynthesis and pathogen defense.

The stomatal density and maximum pore area set an anatomical upper limit on stomatal conductance (Brown and Escombe, 1900; Parlange and Waggoner, 1970; Franks and Farquhar, 2001; Franks and Beerling, 2009b; Lehmann and Or, 2015; Sack and Buckley, 2016; Harrison et al., 2019), but stomatal shape, distribution, and patterning also affect gas exchange. Smaller guard cells and dumbbell-shaped stomata of grasses can respond faster to environmental changes (Drake et al., 2013), but responsiveness is further modulated by subsidiary cell anatomy and physiology (Franks and Farquhar, 2007; Raissig et al., 2017; Gray et al., 2020). Stomatal clustering reduces gas exchange and photosynthesis because adjacent stomata interfere with one another (Dow et al., 2014b), diffusion shells overlap (Lehmann and Or, 2015), and limitations on lateral diffusion of CO_2_ in the mesophyll (Lawson and Blatt, 2014 and references therein). However, sparse clusters of small stomata could allow a leaf with low rates of gas exchange to have faster stomatal response compared to a leaf with large, low-density stomata (Papanatsiou et al., 2017). Leaves with stomata on both lower and upper surfaces (amphistomatic or amphistomatous) supply more CO_2_ to the mesophyll than hypostomatic (aka hypostomatous) leaves that only have stomata on the lower surface (Parkhurst, 1978; Gutschick, 1984; Parkhurst and Mott, 1990; Oguchi et al., 2018). In addition to anatomy, the pore area shrinks and expands in response to internal and external factors to regulate gas exchange dynamically (Buckley, 2019). For example, stomata typically open during the day and close at night in C_3_/C_4_ plants, but the opposite is true for CAM plants. Shade, high vapor pressure deficits, dry soil and other factors can cause stomata to (partially) close even in the middle of the day. Variation in how stomata respond to internal and external signals may explain as much of the variation in gas exchange across leaves as anatomy (Lawson and Blatt, 2014).

Many types of foliar pathogens, including viruses (Murray et al., 2016), bacteria (Melotto et al., 2006; Underwood et al., 2007), protists (Fawke et al., 2015), and fungi (Hoch et al., 1987; Zeng et al., 2010) use stomatal pores to gain entry into the leaf. For example, rust fungi hyphae recognize the angle at which guard cells project from the leaf surface and use it as a cue for appressorium formation (Allen et al., 1991). Oomycete pathogens can target open stomata on a leaf (Kiefer et al., 2002). Plants can limit colonization through innate immunity, called stomatal defense (recently reviewed in Melotto et al., 2017), by closing stomata after they recognize microbe-associated molecular patterns (MAMPs) on pathogen cells. Some bacterial pathogens have responded by evolving the ability to prevent stomatal closure, increasing their colonization of the leaf interior (Melotto et al., 2006). In addition to stomatal closure, anatomical changes in stomatal density and/or size might provide another layer of defense against pathogen colonization. For example, infection increases in leaves with higher stomatal density (McKown et al., 2014; Tateda et al., 2019; Dutton et al., 2019; Fetter et al., 2019). The positive effect of stomatal density on infection suggests that infection is limited by the number or size of locations for colonization, meaning that many individual pathogens must usually be unable to find stomata or other suitable locations for colonization. This is actually somewhat surprising given the ability of some pathogens to search for and sense stomata (see above).

Stomatal anatomy could be a key link between gas exchange and pathogen colonization. Although many anatomical factors and stomatal movement affect gas exchange (see above), here I focus on the density and size of stomata in a hypostomatous leaf. Stomatal size refers to both the area of guard cells when fully open, from which one can calculate the pore area for gas exchange (see Model). For simplicity, I model a hypostomatous leaf, but consider the implications for amphistomatous leaves in the Discussion. Stomatal size and density not only determine the theoretical maximum stomatal conductance (*g*_s,max_), but is also proportional to the operational stomatal conductance (*g*_s,op_) in many circumstances (Franks et al., 2009, 2014; Dow et al., 2014a; McElwain et al., 2016; Murray et al., 2019). *g*_s,op_ is the actual stomatal conductance of plants in the field and is almost always below *g*_s,max_ because stomata are usually not fully open. Although they are not the same, the strong empirical relationship between *g*_s,max_ and *g*_s,op_ means that anatomical *g*_s,max_ can be used as a proxy for *g*_s,op_ without explicitly modeling dynamic changes in stomatal aperture (see Discussion). Stomatal size and density have also been measured on many more species than stomatal responsiveness, which may make it easier to test predictions.

After a pathogen reaches a host, it must survive on the leaf surface and colonize the interior (Beattie and Lindow, 1995; Tucker and Talbot, 2001). For analytical tractability, I restrict the focus here to colonization by a pathogen using a random search on a leaf without stomatal defense (i.e. a leaf that cannot recognize pathogens and close stomata). Obviously, these simplifications ignore a lot of important plant-pathogen interaction biology. In the Discussion, I delve further into these limitations and suggest future work to overcome these limitations. In order for pathogen-mediated selection on stomatal anatomy, I assume that the pathogen reduces host fitness once it colonizes (Gilbert, 2002). Susceptible hosts can lose much of their biomass or die, but even resistant hosts must allocate resources to defense or reduce photosynthesis because of defoliation, biotrophy, or necrosis around sites of infection (Bastiaans, 1991; Mitchell, 2003).

The purpose of this study is to develop a theoretical framework to test whether variation in stomatal size and density arises from a tradeoff between gas exchange and pathogen colonization. Since stomatal size and density affect both gas exchange and pathogen colonization, selection to balance these competing demands could shape stomatal size-density scaling relationships. Botanists have long recognized that stomatal size and density are inversely correlated (Weiss, 1865; Tichá, 1982; Hetherington and Woodward, 2003; Sack et al., 2003; Franks and Beerling, 2009a; Brodribb et al., 2013; Boer et al., 2016), but the evolutionary origin of this relationship is not yet known. Here I argue that deleterious effects of pathogen infection could shape selection on this relationship. Explanations for inverse size-density scaling are usually cast in terms of preserving *g*_s,max_ and/or stomatal cover (*f*_S_), defined at the fraction of epidermal area allocated to stomata (Boer et al., 2016), because there are many combinations of stomatal size and density that have same *g*_s,max_ or same *f*_S_:

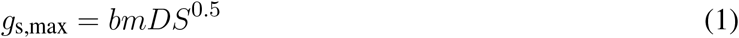

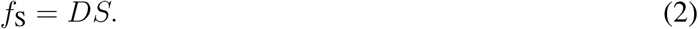

*D* and *S* are stomatal stomatal density and size, respectively (see Table 1 for a glossary of mathematical symbols and units). *b* and *m* are assumed to be biophysical and morphological constants, *sensu* Sack and Buckley (2016) (see Supplementary Material). *f*_S_ is proportional to the more widely used stomatal pore area index [Sack et al. (2003); see Supplementary Material]. If size and density also affect pathogen colonization, then selection from foliar pathogens could significantly alter the size-density scaling relationship. The empirical size-density scaling relationship is linear on a log-log scale, determined by an intercept *α* and slope *β*:

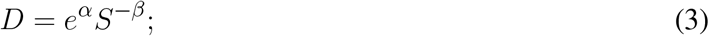

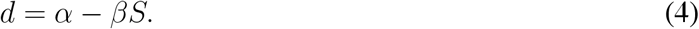

**Table 1.**
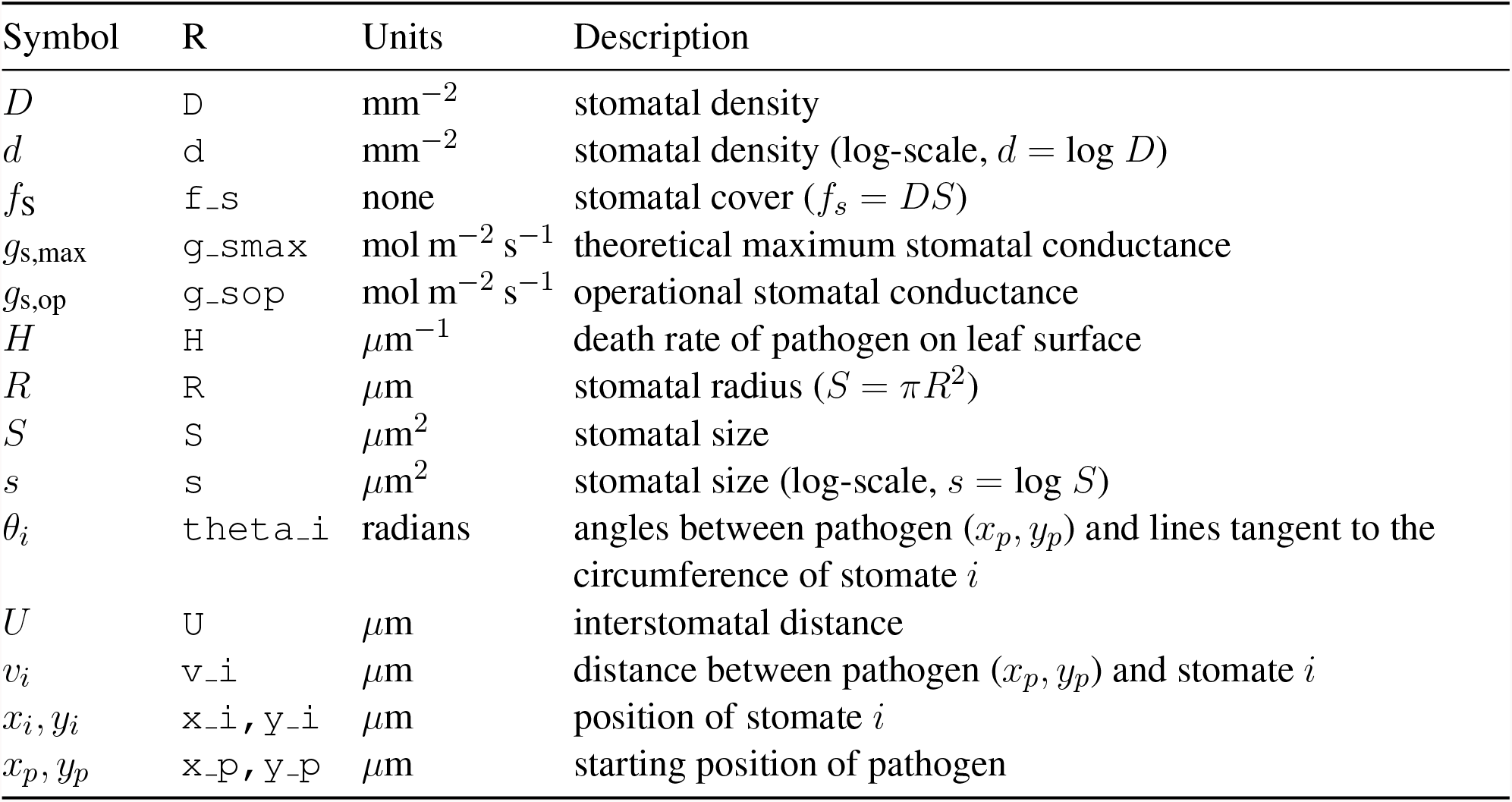
Glossary of mathematical symbols. The columns indicate the mathematical Symbol used in the paper, the associated symbol used in R scripts, scientific Units, and a verbal Description.

| Symbol | R | Units | Description |
| --- | --- | --- | --- |
| $D$ | D | $\text{mm}^{-2}$ | stomatal density |
| $d$ | d | $\text{mm}^{-2}$ | stomatal density (log-scale, $d = \log D$) |
| $f_s$ | f_s | none | stomatal cover ($f_s = DS$) |
| $g_{s,\max}$ | g_smax | $\text{mol m}^{-2} \text{s}^{-1}$ | theoretical maximum stomatal conductance |
| $g_{s,\text{op}}$ | g_sop | $\text{mol m}^{-2} \text{s}^{-1}$ | operational stomatal conductance |
| $H$ | H | $\mu\text{m}^{-1}$ | death rate of pathogen on leaf surface |
| $R$ | R | $\mu\text{m}$ | stomatal radius ($S = \pi R^2$) |
| $S$ | S | $\mu\text{m}^2$ | stomatal size |
| $s$ | s | $\mu\text{m}^2$ | stomatal size (log-scale, $s = \log S$) |
| $\theta_i$ | theta_i | radians | angles between pathogen ($x_p, y_p$) and lines tangent to the circumference of stomate $i$ |
| $U$ | U | $\mu\text{m}$ | interstomatal distance |
| $v_i$ | v_i | $\mu\text{m}$ | distance between pathogen ($x_p, y_p$) and stomate $i$ |
| $x_i, y_i$ | x_i, y_i | $\mu\text{m}$ | position of stomate $i$ |
| $x_p, y_p$ | x_p, y_p | $\mu\text{m}$ | starting position of pathogen |

For brevity, *d* = log(*D*) and *s* = log(*S*). Rearranging Equations 1 and 2, a scaling relationship where *β* = 0.5 preserves *g*_s,max_ while *β* = 1 preserves *f*_S_.

How would adding pathogens alter these predicted scaling relationships? For simplicity, consider two environments, one without foliar pathogens and one with lots. In the absence of foliar pathogens, we expect size-density scaling to preserve *g*_s,max_, *f*_S_, or some least-cost combination of them. What happens when we introduce pathogens? If stomatal size and density increase pathogen colonization, then selection will favor reduced size and/or density. This would change the intercept *α* but not the slope. The effect of foliar pathogens on the slope depends on the relationship between size, density, and probability of colonization. If the probability of colonization is proportional to the product of *linear* stomatal size (*S*^0.5^) and density (∝ *DS*^0.5^ as for *g*_s,max_) then it has the same effect on the slope as *g*_s,max_ because there are many combinations of *D* and *S*^0.5^ that have same probability of colonization. If the probability of colonization is proportional to the product of *areal* stomatal size (*S*) and density (∝ *DS* as for *f*_S_) then it has the same effect on the slope as *f*_S_ because there are many combinations of *D* and *S* that have same probability of colonization. Alternatively, the probability of colonization may have a different scaling relationship (neither 0.5 nor 1) or may be nonlinear on a log-log scale. Unlike *g*_s,max_ and *f*_S_, we do not have theory to predict a stomatal size-density relationship that preserves the probability of colonization.

In summary, the physical relationship between stomatal size, density, and conductance is well established (Harrison et al., 2019). Size and density also likely affect the probability of pathogen colonization, but we do not have a theoretical model that makes quantitative predictions. The inverse stomatal size-density relationship has usually been explained in terms of preserving stomatal conductance and/or stomatal cover, but selection by pathogens might alter scaling. To address these gaps, the goals of this study are to 1) introduce a spatially explicit model pathogen colonization on the leaf surface; 2) use the model to predict the relationship between *g*_s,max_, *f*_S_, and the probability of colonization; 3) work out what these relationships predict about stomatal size-density scaling. I analyzed an idealized, spatially explicit Model of how a pathogen lands on a leaf and finds a stomate to colonize the leaf using a random search. To my knowledge, this is the first model that makes quantitative predictions about the relationship between stomatal anatomy, the probability of colonization, and their impact on stomatal size-density scaling.

## MODEL

In this section, I introduce a spatially explicit model of successful pathogen colonization on a leaf surface. I explain the model structure and assumptions here; the Materials and Methods section below describes how I analyzed the model to address the goals of the study. For generality, I refer to a generic “pathogen” that lands on leaf and moves to a stomate. The model is agnostic to the type of pathogen (virus, bacterium, fungus, etc.) and the specific biological details of how it moves. For example, motile bacterial cells can land and move around (Beattie and Lindow, 1995) whereas fungi may germinate from a cyst and grow until they form an appresorium for infection (Tucker and Talbot, 2001). These very different tropic movements on the leaf are treated identically here. I do not model photosynthesis explicitly, but assume that stomatal conductance limits carbon fixation, even though the relationship is nonlinear. I used Sympy version 1.6.1 (Meurer et al., 2017) for symbolic derivations.

### Spatial representation of stomata

Stomata develop relatively equal spacing to minimize resistance to lateral diffusion (Morison et al., 2005), allow space between stomata (Dow et al., 2014b), and prevent stomatal interference (Lehmann and Or, 2015). Here I assume that stomata are arrayed in an equilateral triangular grid with a density *D* and size (area) *S* on the abaxial surface only, since most leaves are hypostomatoues [Muir (2015); but see Discussion]. This assumption ignores veins, trichomes, and within-leaf variation in stomatal density. Stomata are therefore arrayed in an evenly spaced grid (Figure 1a). The interstomatal distance *U*, measured as the distance from the center of one stomata to the next, is the maximal diagonal of the hexagon in *µ*m that forms an equal area boundary between neighboring stomata. The area of a hexagon is 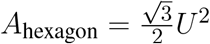. By definition the stomatal density is the inverse of this area, such that 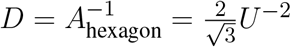. Therefore, interstomatal distance can be derived from the stomatal density as:

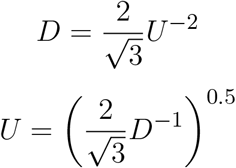

**Figure 1.**
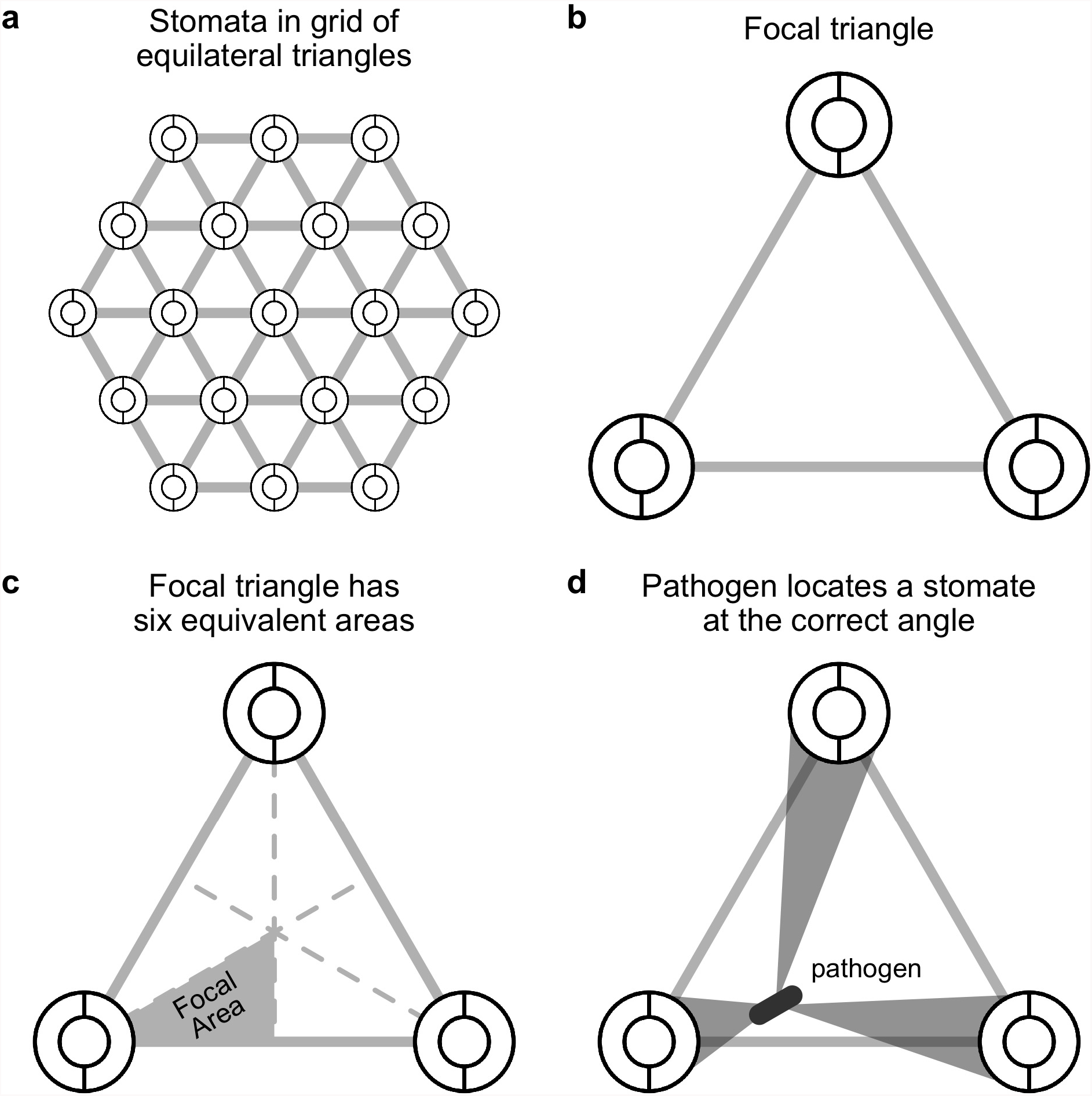
A spatially explicit model of stomatal anatomy and pathogen colonization. **a**. Stomata are assumed to be in a homogenous equilateral triangular grid, which means that we can extrapolate from **b.** a focal triangle to the entire leaf. The circles represent idealized stomata; the grey lines between them are for visualization. **c.** By symmetry, a single focal region within the focal triangle can be modeled and extrapolated to the rest of the triangle. **d.** The model assumes that a pathogen, depicted as a grey rod, lands somewhere on the leaf surface and will sucessfully locate a stomate if it moves at the correct angle, depicted by the grey polygons.

For example, if the density is *D* = 10^2^ mm^−2^ = 10^−4^ *µ*m^−2^, then *U* is 107.5 *µ*m. Parkhurst (1994) described this result previously. I also make the simplifying assumption that stomata are perfectly circular with radius *R* when fully open. This may be approximately true for fully open stomata with kidney-shaped guard cells (Sack and Buckley, 2016 and references therein). Although I assume stomata are circular here, in calculating *g*_s,max_, I assume typical allometric relationships between length, width, and pore area [Sack and Buckley (2016); see Supplementary Material].

### Spatial representation of pathogen search

Since stomata are arrayed in a homogeneous grid, we can focus on single focal triangle (Figure 1b-c). Suppose that an individual pathogen (e.g. bacterial cell or fungal spore) lands at a uniform random position within the focal triangle and must arrive at a stomate to colonize. If it lands on a stomate, then it infects the leaf with probability 1; if it lands between stomata, then it infects the leaf with probability *p*_locate_. This is the probability that it locates a stomate, which I will derive below. The probabilities of landing on or between a stomate are *f*_S_ and 1 *− f_S_*, respectively. Hence, the total probability of colonization is:

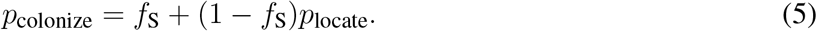

I assume that the pathogen cannot sense where stomata are and orients at random, thereafter traveling in that direction. If it successfully locates a stomate, it colonizes the leaf, but otherwise does not infect. If there is a high density of stomata and/or large stomata, the probability of locating a stomate increases. By assuming that stomata form an equilateral triangular grid (see above), we can extrapolate what happens in the focal triangle (Figure 1b) by symmetry. Further, since an equilateral triangle can be broken up into six identical units (Figure 1c), we can simply calculate *p*_locate_ in this focal area. This implicitly assumes that the probability of colonizing stomata outside the focal area is 0 because they are too far away. This assumption may be unrealistic for larger pathogens, such as fungi, whose hyphae can travel longer distances on the leaf surface (Brand and Gow, 2012). In Appendix 1: Spatially implicit model I derive a simpler, but spatially *implicit* model that relaxes the assumption the pathogens must colonize a stomate within their focal triangle.

Consider a pathogen that lands in position (*x_p_, y_p_*) within the triangle. The centroid of the triangle is at position (*x_c_, y_c_*) and a reference stomate is at position (0, 0) (Figure 2a). Therefore *x_c_* = *U*/2 and 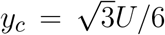. The other stomata are at positions 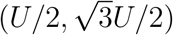 and (*U,* 0) (Figure 2). *x_p_* and *y_p_* are defined as the horizontal and vertical distances, respectively, from the pathogen to the reference stomate at position (0, 0).

**Figure 2.**
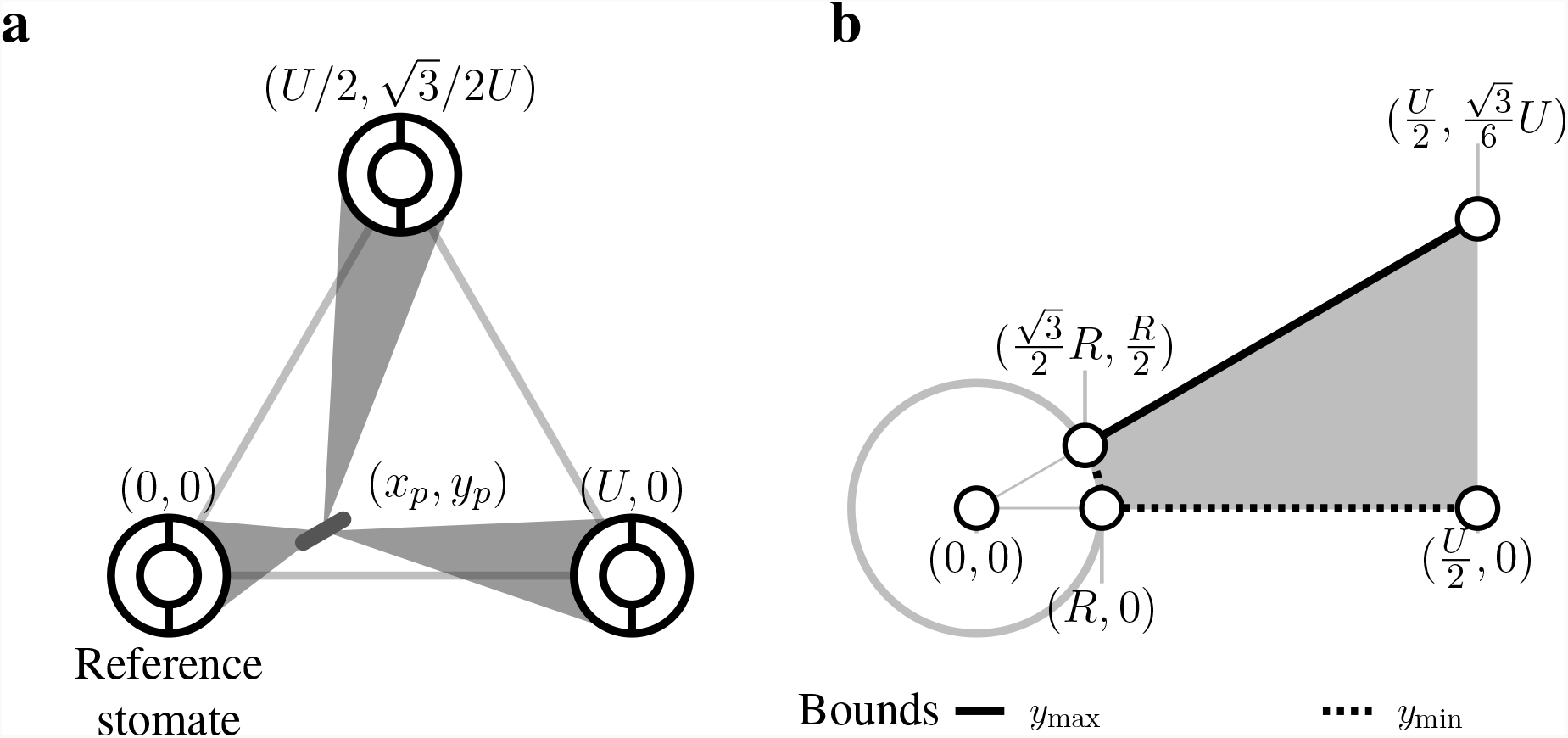
Spatial representation of stomata and pathogen. **a.** The pathogen starts at a uniform random position within the focal region denoted (*x_p_, y_p_*). Within the focal triangle, the reference stomate is at position (0,0) by definition, and other stomatal positions are determined by the interstomatal distance *U*. **b.** Within the focal region, a pathogen can land within the stomate (white circle with grey outline and radius *R*) or in the grey area. The outer borders of this area are shown and depend on *R* and *U*. For a given position *x*, there is a minimum *y*-value (*y*_min_, dashed line) and maximum *y*-value (*y*_max_, solid line).

Given that the pathogen starts at position (*x_p_, y_p_*), what’s the probability of contacting one of the stomata at the vertices of the focal triangle? I assume the probability of contacting a stomate is equal to the proportion of angular directions that lead to a stomate (Figure 1d). I solved this by finding the angles (*θ*_1_, *θ*_2_, *θ*_3_) between lines that are tangent to the outside of the three stomata and pass through (*x_p_, y_p_*) (Figure 2a). If stomate *i* is centered at (*x_i_, y_i_*), the two slopes of tangency as function of pathogen position are:

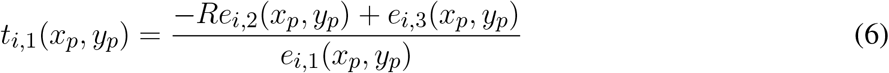

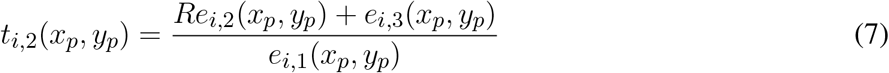

where

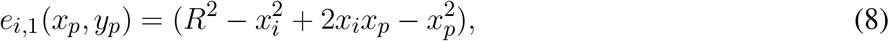

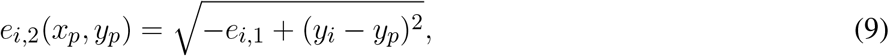

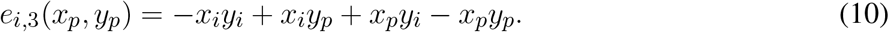

Note that *i* ∈ {1, 2, 3}, indexing the three stomata in the focal triangle. The angle in radians between *t*_*i*,1_(*x_p_, y_p_*) and *t_i,_*_2_(*x_p_, y_p_*) is:

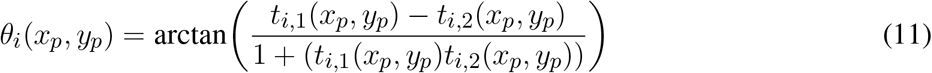

I further assumed that the longer distance a pathogen must travel, the less likely it would be to locate a stomate. For example, if stomata are at very low density, then a pathogen may die before it reaches a stomate because of UV, desiccation, or another factor. I included this effect by assuming the probability of reaching a stomate declines exponentially at rate *H* with the Euclidean distance *v_i_*(*x_p_, y_p_*) between the pathogen location and the edge of stomata *i*, which is distance *R* from its center at *x_i_, y_i_*:

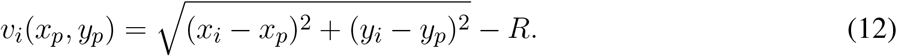

The probability of locating a stomate as a function of pathogen position (*x_p_* and *y_p_*) is the sum of the angles divided by 2*π*, discounted by their distance from the stomate:

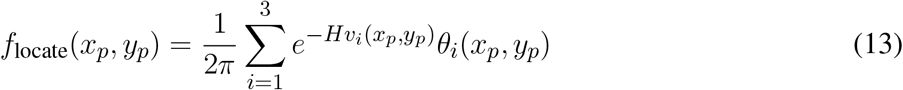

When there is no pathogen death (*H* = 0), *p*_locate_ is the fraction of angles that lead from (*x_p_, y_p_*) to a stomate. When *H* > 0, *p*_locate_ is proportional to this fraction, but less than it depending on stomatal density, size, and starting location of the pathogen.

To obtain the average *p*_locate_, we must integrate *f*_locate_(*x_p_, y_p_*) over all possible starting positions (*x_p_, y_p_*) within the focal area. The focal area is a 30-60-90 triangle with vertices at the center of the reference stomate (0, 0), the midpoint of baseline (*U*/2, 0), and the centroid of the focal triangle 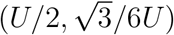 (Figure 1c). Colonization occurs with probability 1 if the pathogen lands in the reference stomate, so we need to integrate the probability of colonization if it lands elsewhere. This region extends from the edge of the stomate, at 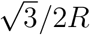 to *U*/2 (Figure 2b). At any *x*, we integrate from the bottom of the focal area (*y*_min_) to the top (*y*_max_):

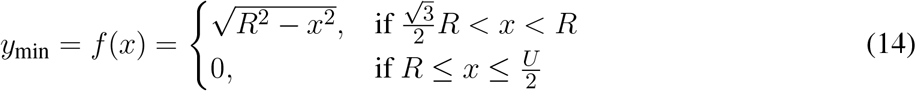

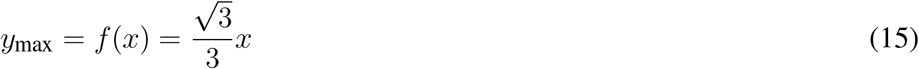

The integral is:

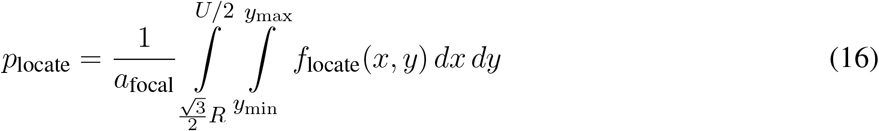

*a*_focal_ is the area of the focal region depicted in grey in Figure 2b:

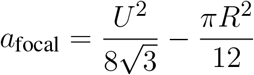

## MATERIALS AND METHODS

The Model calculates a probability of host colonization (Equation 5) as a function of stomatal density, size, and position of a pathogen on the leaf. I solved *p*_colonize_ by importing symbolic derivations from Sympy into R with **reticulate** version 1.16 (Ushey et al., 2020) and used the `integral2()` function in the **pracma** package version 2.2.9 (Borchers, 2019) for numerical integration. I used R version 4.0.2 (R Core Team, 2020) for all analyses and wrote the paper in **rmarkdown** version 2.3 (Xie et al., 2018; Allaire et al., 2020). Citations for additional R software packages are in Appendix 2. Source code is deposited on GitHub (https://github.com/cdmuir/stomata-tradeoff) and will be archived on Zenodo upon publication.

### What is the relationship between stomatal size, density, and colonization?

I calculated *p*_colonize_ over a biologically plausible grid of stomatal size and density for hypostomatous species based on Boer et al. (2016). Stomatal density (*D*) ranges from 10^1^ − 10^3.5^ mm^−2^; stomatal size (*S*) ranges from 10^1^ − 10^3.5^ *µ*m^2^. I only considered combinations of size and density where stomatal cover (*f*_S_) was less than 1/3, which is close to the upper limit in terrestrial plants (Boer et al., 2016). I crossed stomatal traits with three levels of *H* ∈ {0, 0.01, 0.1}. When *H* = 0, a pathogen persists indefinitely on the leaf surface. *H* = 0.01 and *H* = 0.1 correspond to low and high death rates, respectively. These values are not necessarily realistic, but illustrate qualitatively how a hostile environment on the leaf surface alters model predictions.

### How do pathogens alter optimal stomatal size-density scaling?

The stomatal size-density scaling relationship can be explained in terms of preserving a constant stomatal conductance (*g*_s,max_) that is proportional to *DS*^0.5^ when *bm* is constant (Equation 1). In other words, there are infinitely many combinations of *D* and *S*^0.5^ with the same *g*_s,max_. If *g*_s,max_ is held constant at *C_g_*, then the resulting size-density scaling relationship on a log-log scale is:

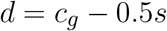

where lowercase variables are log-transformed equivalents of their uppercase counterparts (Table 1). The scaling exponent *β_g_* = 0.5 preserves *C_g_*.

Next, suppose there is a scaling exponent *β_p_* that preserves *p*_colonize_ for the product *DS^βp^*. If *β_p_* = 0.5, then *p*_colonize_ is always proportional to *g*_s,max_. If *β_p_* > 0.5, small, densely packed stomata would be more resistant to colonization (lower *p*_colonize_) compared to larger, sparsely spaced stomata with the same *g*_s,max_. If *β_p_* < 0.5, small, densely packed stomata would be less defended (higher *p*_colonize_) compared to larger, sparsely spaced stomata with the same *g*_s,max_. I refer to the three outcomes (*β_p_* = 0.5, *β_p_* < 0.5, and *β_p_* > 0.5) as iso-, hypo-, and hyper-conductance, respectively. I was unable to solve analytically for *β_p_*, so I numerically calculated isoclines of *p*_colonize_ over the grid of *D* and *S* values described in the preceding subsection. I numerically calculated the scaling relationships at a constant *p*_colonize_ ∈ {0.025, 0.1, 0.4} for *H* ∈ {0, 0.01, 0.1}.

## RESULTS

### Nonlinear relationships between colonization, stomatal cover, and conductance

The probability of colonization (*p*_colonize_) is not simply proportional to stomatal cover (*f*_S_). At low *f*_S_, *p*_colonize_ increases rapidly relative to *f*_S_ at first Figure 3a). At higher *f*_S_, *p*_colonize_ increases linearly with *f*_S_. When pathogens persist indefinitely (*H* = 0), any combination of stomatal size (*S*) and density (*D*) with the same *f*_S_ have the same effect on *p*_colonize_. When *H* > 0, pathogens are less likely to land close enough to a stomate to infect before dying, so *p*_colonize_ is closer to *f*_S_ (Figure 3a). The maximum *p*_colonize_ under the range of parameters considered was ~ 0.6 when *H* = 0 and *f*_S_ is at its maximum value of 1/3. When *f*_S_ is low, *p*_colonize_ is also low. The relationship between *p*_colonize_, *f*_S_, and *g*_s,max_is qualitatively similar in the spatially implicit model, but the values for *p*_colonize_ are substantially higher because pathogens can potentially colonize any stomate on the leaf rather than only those in the focal triangle (see Appendix 1: Spatially implicit model for more detail). Bear in mind that this is the probability for a single individual searching randomly; if enough individuals reach the leaf and/or they can actively find stomata, it’s almost certain that at least some will colonize the leaf. However, reducing *p*_colonize_ may help plants limit the damage since fewer total individual pathogens will colonize the leaf interior.

**Figure 3.**
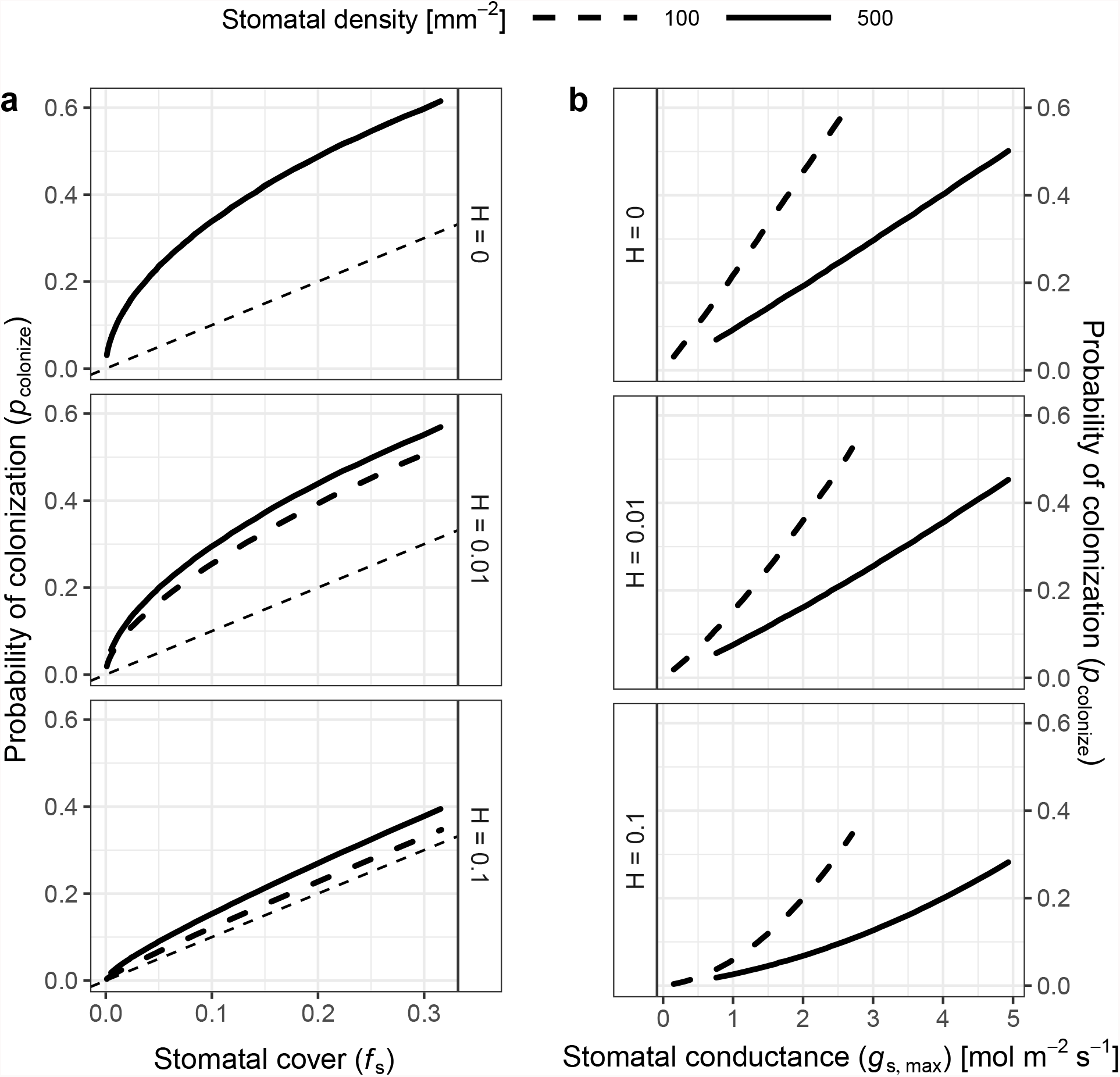
The probability of colonization increases with both stomatal cover and conductance. I simulated the probability of colonization (*p*_colonize_, *y*-axis) over a range of stomatal densities and sizes (see [Materials and Methods]), but a subset of results are shown here. Stomatal size and density determine stomatal cover (*f*_S_; Equation 2) and theoretical maximum stomatal conductance (*g*_s,max_; Equation 1). **a.** *p*_colonize_ initially increases rapidly with *f*_S_ (*x*-axis), then slows down to a linear relationship. Overall, *p*_colonize_ is lower when pathogens can die on the leaf surface (*H* > 0). The relationship between *f*_S_ and *p*_colonize_ is the same regardless of stomatal density when *H* = 0 (upper facet), which is why the lines overlap. When *H* > 0, higher density (solid lines) increase *p*_colonize_ (lower facets). **b.** *p*_colonize_ increases exponentially with *g*_s,max_ at all stomatal densities, but *p*_colonize_ is much lower at higher densities for a given *g*_s,max_. The relationship between *g*_s,max_ and *p*_colonize_ is similar for all values of *H*.

*p*_colonize_ is not directly proportional to *f*_S_ because it depends on *D* and *S* in quantitatively different ways (Figure S1). For the same *f*_S_, leaves with greater *D* have higher *p*_colonize_ (Figure 3a). Holding *f*_S_ constant, leaves with lower *D* and higher *S* will have a greater distance (*v_i_*) between a pathogen and its stomata. When *H* > 0, this extra distance leads more pathogens to die before they can find a stomate. However, this result is inconsistent with the spatially implicit model (Appendix 1) because *S* and *D* have identical effects on *f*_S_.

In contrast to *f*_S_, *p*_colonize_ increases at a greater than linear rate with stomatal conductance (*g*_s,max_). Greater *D* (smaller *S*) is associated with lower *p*_colonize_ for a given value of *g*_s,max_ (Figure 3b). This happens because *p*_colonize_ increases approximately linearly with *S* whereas *g*_s,max_ is proportional to *S*^0.5^. Therefore, *p*_colonize_ increases exponentially with *g*_s,max_ at all stomatal densities, but the rate of growth is lower at greater *D* for a given value of *g*_s,max_.

### Hyper-conductance size-density scaling

The scaling relationship between *S* and *D* that preserves *p*_colonize_ is always greater than 0.5 (hyper-conductance), but usually less than 1. When *H* = 0, the scaling relationship is essentially 1 (Figure 4), which means that an increase *f*_S_ leads to a proportional increase in *p*_colonize_. Because the scaling relationship is greater than 0.5, leaves with greater stomatal density will have lower *p*_colonize_ than leaves lower stomatal density but the same *g*_s,max_. In other words, increasing *D* and lowering *S* allows plants to reduce *p*_colonize_ while maintaining *g*_s,max_. The scaling relationship is slightly less than 1, but still greater than 0.5, when *H* > 0 (Figure 4). In this area of parameter space, lower stomatal density can reduce *f*_S_ while *p*_colonize_ is constant, but this will still result in lower *g*_s,max_. In the spatially implicit model, the size-density scaling exponent was always exactly 1 except when *H* = 0 (Appendix 1).

**Figure 4.**
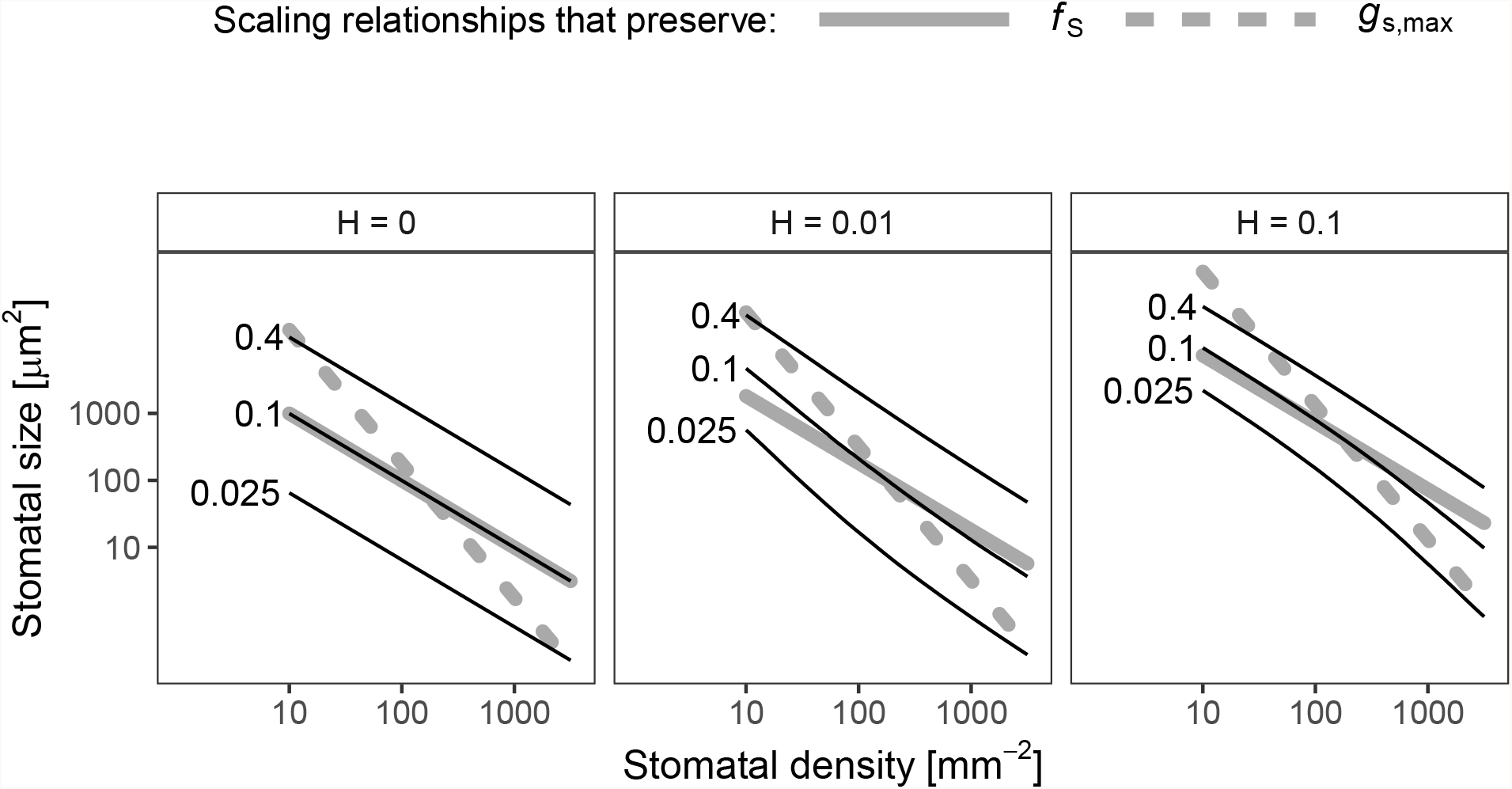
Log-log scaling relationships between stomatal density (*D*,*x*-axis) and size (*S*,*y*-axis) that preserve the probability of colonization (*p*_colonize_). In each panel, solid lines indicate values of *D* and *S* where *p*_colonize_ is 0.025 (lowest line), 0.1, or 0.4 (highest line). For reference, dashed grey lines show scaling relationships that preserve *f*_S_ (*β* = 1, slope = 1/*β* = 1) and *g*_s,max_ (*β* = 0.5, slope = 1/*β* = 2) drawn through the centroid of the plotting region. When the death rate on the leaf surface is low (*H* = 0), the scaling exponent is very close to *β* = 1. When *H* > 0, 0.5 *< β* < 1 and is slightly nonlinear on a log-log scale.

## DISCUSSION

Stomatal density and size set the upper limit on gas exchange in leaves (Harrison et al., 2019) and is often closely related to operational stomatal conductance in nature (Murray et al., 2019). Despite the fact that many foliar pathogens infect through stomata, the relationship between stomatal anatomy and resistance to foliar pathogens is less clear than it is for gas exchange. I used a spatially explicit model of a pathogen searching for a stomate to colonize a host. From this Model, I derived predictions about the relationship between stomatal anatomy and the probability of colonization, a component of disease resistance. The model predicts that the probability of colonization is not always proportional to the surface area of leaf covered by stomata (*f*_S_), as one might intuitively predict. If the leaf surface is a hostile environment and pathogens have a limited time to search, lower stomatal density decreases the probability of colonization even if *f*_S_ is constant. However, *g*_s,max_ decreases proportionally more than the probability of colonization. The model highlights the potential for conflicting demands of minimizing pathogen colonization, minimizing stomatal cover, and maintaining stomatal conductance. Including the effect of anatomy on pathogen colonization therefore has the potential to change our understanding of how stomatal size-density scaling evolves in land plants.

The model predicts that in most cases, increasing stomatal cover should lead to a proportional increase in colonization, which agrees with empirical studies (e.g. McKown et al. (2014); Tateda et al. (2019); Dutton et al. (2019); Fetter et al. (2019)). It also makes new, testable predictions that are less intuitive (Table 2). At very low *f*_S_, there is a rapid increase in colonization (Figure 3a). If there are no stomata, the probability of colonization is 0, so the first few stomata dramatically increase the probability. This is less likely to be significant for abaxial (lower) leaf surfaces, which usually have most of the stomata (Salisbury, 1928; Metcalfe and Chalk, 1950; Mott et al., 1984; Peat and Fitter, 1994; Jordan et al., 2014; Muir, 2015; Bucher et al., 2017; Drake et al., 2019). However, many adaxial (upper) leaf surfaces have zero or very few stomata. Using adaxial leaf surfaces, it should be possible to test if small changes in stomatal size or density have a larger effect on pathogen colonization when *f*_S_ is low. Such experiments could use natural genetic variation (McKown et al., 2014) or mutant lines (Dow et al., 2014b). The nonlinear increase in *p*_colonize_ is less apparent when *H* > 0 (Figure 3a). A more hostile microenvironment (e.g. drier, higher UV) should therefore reduce the effect of increased size or density at low *f*_S_. If true, the diminishing marginal effect of *f*_S_ on colonization could explain why stomatal ratio on the upper and lower surface is bimodal (Muir, 2015). The initial cost of adaxial (upper) stomata is relatively high, but if the benefits outweigh the costs, then equal stomatal densities on each surface maximize CO_2_ supply for photosynthesis (Parkhurst, 1978; Gutschick, 1984; Parkhurst and Mott, 1990). The costs and benefits will certainly vary with environmental conditions as well. Future work should extend this model, which considered hypostomatous leaves, to address stomatal size and density in amphistomatous leaves, since leaf surfaces may differ in the type of pathogens present and microenvironment (McKown et al., 2014; Fetter et al., 2019).

**Table 2.** New testable model predictions and suggested experiments to test them.

| Model Prediction | How to test it |
| --- | --- |
| Increasing stomatal size and/or density will have a larger effect on pathogen colonization in leaves with low stomatal cover. | Compare the effect of changing stomatal size and/or density on pathogen colonization in leaves with low and high stomatal cover. |
| Increasing stomatal size and/or density will have a smaller effect on pathogen colonization in epiphytic environments more hostile to pathogens. | Compare the effect of changing stomatal size and/or density on pathogen colonization in more hostile environments (e.g. drier, higher light) |
| When selection against pathogen colonization stronger, the stomatal size-density scaling exponent should be lower | Measure stomatal anatomy in environments that differ in pathogen colonization using comparative or experimental approaches |

An effect of stomatal size and density on foliar pathogen colonization could change our understanding of stomatal size-density scaling. Since allocating leaf epidermis to stomata may be costly (Franks and Farquhar, 2007; Assmann and Zeiger, 1987; Dow et al., 2014b; Lehmann and Or, 2015; Baresch et al., 2019), selection should favor leaves that achieve a desired *g*_s,max_ while minimizing *f*_S_ (Boer et al., 2016). Because of their different scaling exponents (Equation 1, 2), smaller, densely packed stomata can achieve the same *g*_s,max_ at minimum *f*_S_. However, many leaves have larger, sparsely packed stomata. Incorporating pathogen colonization may explain why. If pathogens have a limited time to find stomata before dying (*H* > 0), then the scaling exponent between size and density that keeps *p*_colonize_ constant is between 0.5 and 1, the scaling exponents for *g*_s,max_ and *f*_S_, respectively (Figure 4). Greater density of smaller stomata can increase *g*_s,max_ while keeping *p*_colonize_ constant, but this will increase *f*_S_. Conversely, *f*_S_ could decrease while keeping *p*_colonize_ constant, but this will decrease *g*_s,max_. This sets up the potential for conflict between competing goals. The optimal stomatal size and density will therefore depend on the precise costs and benefits of infection, stomatal conductance, and stomatal cover. This may explain why many leaves have large, sparsely packed stomata despite the fact that they could achieve the same *g*_s,max_ and lower *f*_S_ with smaller, more densely packed stomata.

The model examines the probability of colonization for a single pathogen. The calculated probabilities of colonization should not be interpreted as exact predictions, but rather as depicting qualitative relationships between stomatal anatomy and infection severity. The energetic cost and lost photosynthetic capacity (closed stomata, necrosis, etc.) of dealing with a pathogen is assumed to be proportional to the amount of infection. The actual fitness cost will be modulated by the number of pathogens landing on the leaf and the cost of infection, all else being equal. In environments with fewer or less virulent pathogens, the fitness cost of infection will be less than in environments with more abundant, virulent pathogens. The model is less relevant to very susceptible host plants that can be severely damaged or killed by a small number of colonizations that spread unchecked throughout the host tissue.

## CONCLUSION

The model makes two non-intuitive predictions. First, the effect of increased stomatal density or size on susceptibility to foliar pathogens is greatest when stomatal cover is very low. Second, maximizing disease resistance sets up a potential conflict between minimizing stomatal cover and maximizing stomatal conductance. The first prediction is consistent with results in *Populus trichocarpa* (McKown et al., 2014) and may be relatively straightforward to test experimentally with adaxial (upper) stomata that occur at low and moderate densities within the same or closely related species (Muir et al., 2014; Fetter et al., 2019). The second prediction about size-density scaling is more complex because we would need to know the relationships between colonization, stomatal cover, stomatal conductance, and fitness in natural conditions. There is growing evidence that stomata mediate tradeoffs between photosynthesis and defense in *Populus trichocarpa* (McKown et al., 2019), but testing these predictions in a variety of species will help determine whether pathogens have played an important role shaping stomatal anatomy in land plants.

## ACKNOWLEDGMENTS

I would like to thank Athena McKown and an anonymous referee for valuable feedback that improved this manuscript.

## SUPPLEMENTARY MATERIAL

### *g*_s,max_ calculation

I calculated *g*_s,max_ (Equation 1) to water vapor at a reference leaf temperature (*T*_leaf_ = 25° C) following Sack and Buckley (2016). They defined a biophysical and morphological constant as:

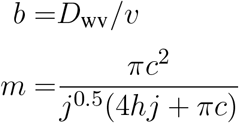

*b* is the diffusion coefficient of water vapor in air (*D*_wv_) divided by the kinematic viscosity of dry air (*v*). *D*_wv_ = 2.49 × 10^−5^ m^2^ s^−1^ and *v* = 2.24 × 10^−2^ m^3^ mol^−1^ at 25° (Monteith and Unsworth, 2013). For kidney-shaped guard cells, *c* = *h* = *j* = 0.5.

### *f*_S_ is proportional to the stomatal pore area index

The stomatal pore area index (SPI; Sack et al., 2003) is calculated as the product of the stomatal density and guard cell length (GCL) squared:

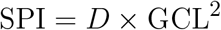

Assuming that the stomatal radius *R* is half the GCL, then stomatal size *S* is equivalent to:

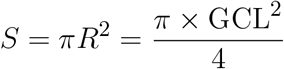

Based on equation 2, it follows that *f*_S_ and SPI are proportional:

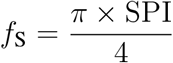

---

**Figure S1.**
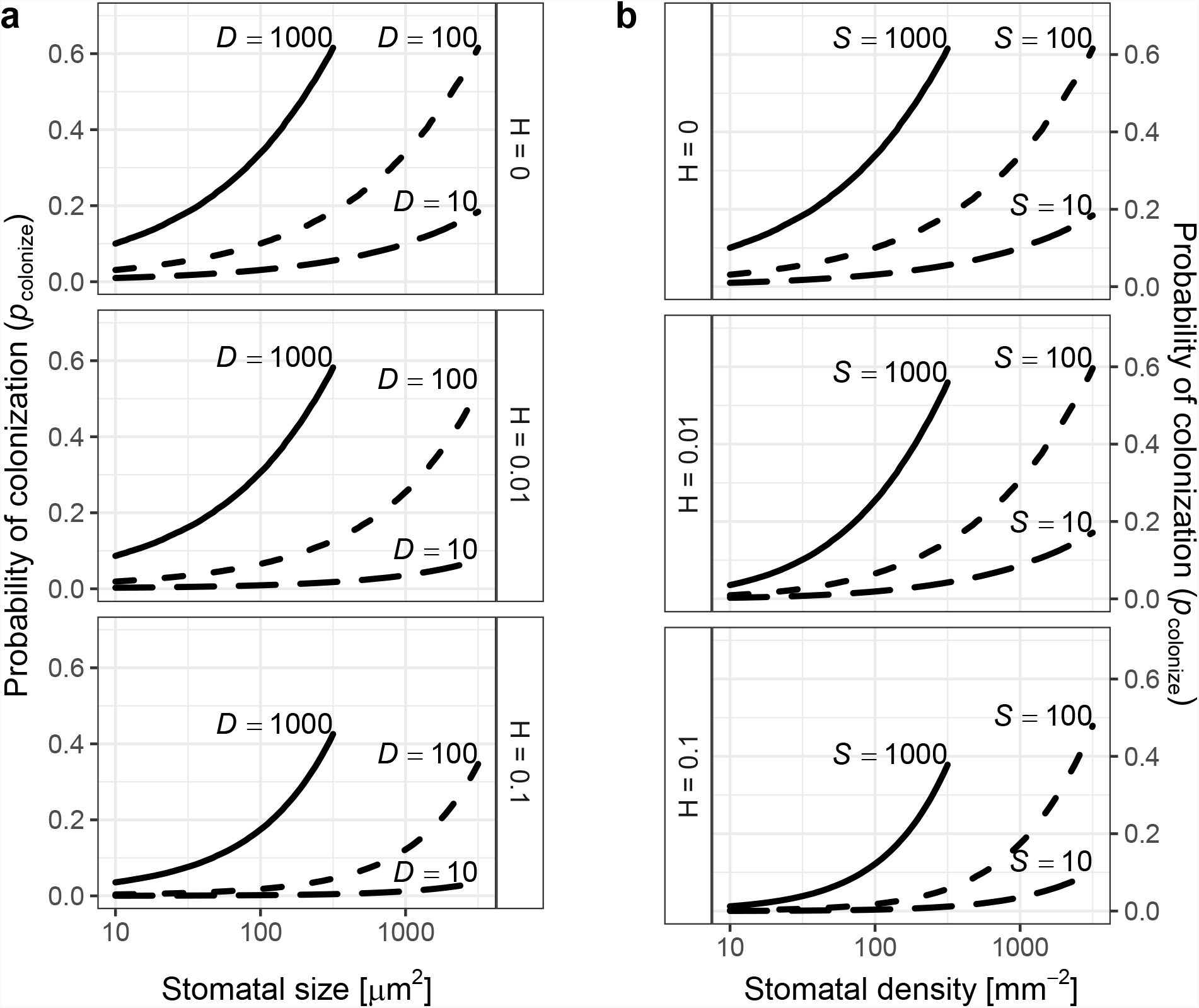
The probability of colonization increases with both stomatal size (*S*) and density *D*. I simulated the probability of colonization (*p*_colonize_, *y*-axis) over a range of *S*, *D*, and *H* (see [Materials and Methods]) **a.** Each line shows how *p*_colonize_ increases with *S* (*x*-axis, log-scale) for selected values of *D* ∈ {10, 100, 1000} mm^−2^. **b.** Each line shows how *p*_colonize_ increases with *D* (*x*-axis, log-scale) for selected values of *S* ∈ {10, 100, 1000} *µ*m^2^. The facets show results for different values of *H*

## Figures

**Figure S2.**
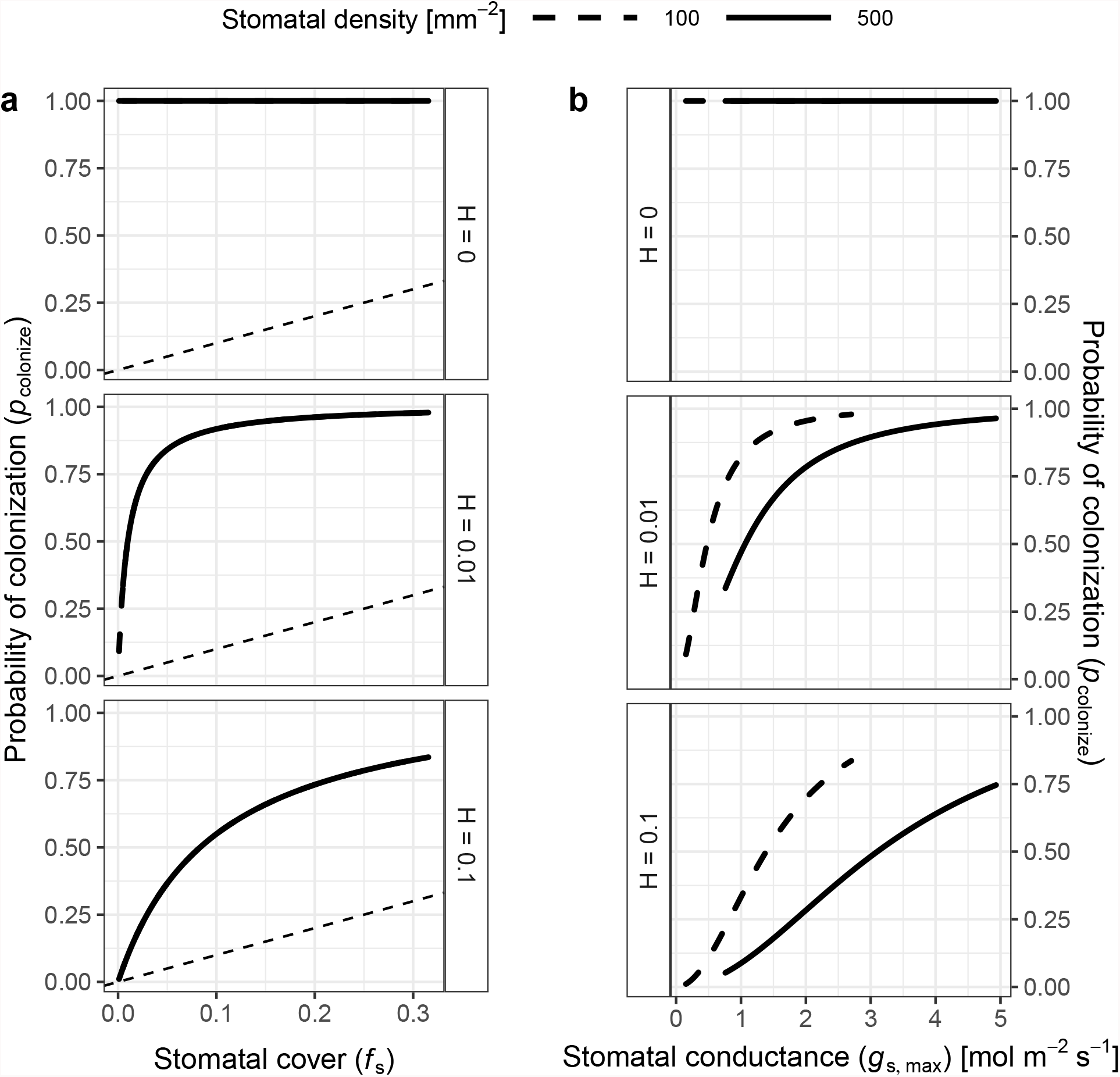
The probability of colonization increases with both stomatal cover and conductance in the spatially implicit model. As in Figure 3, I simulated the probability of colonization (*p*_colonize_, *y*-axis) over a range of stomatal densities and sizes (see [Materials and Methods]), but a subset of results are shown here. Stomatal size and density determine stomatal cover (*f*_S_; Equation 2) and theoretical maximum stomatal conductance (*g*_s,max_; Equation 1). **a.** *p*_colonize_ initially increases rapidly with *f*_S_ (*x*-axis), then slows down to a linear relationship. Overall, *p*_colonize_ is lower when pathogens can die on the leaf surface (*H* > 0). The relationship between *f*_S_ and *p*_colonize_ is the same regardless of stomatal density for all values of *H*, which is why the lines overlap. **b.** *p*_colonize_ increases sigmoidally with *g*_s,max_ at all stomatal densities, but *p*_colonize_ is lower at higher densities for a given *g*_s,max_. The relationship between *g*_s,max_ and *p*_colonize_ is similar for all values of *H* > 0.

**Figure S3.**
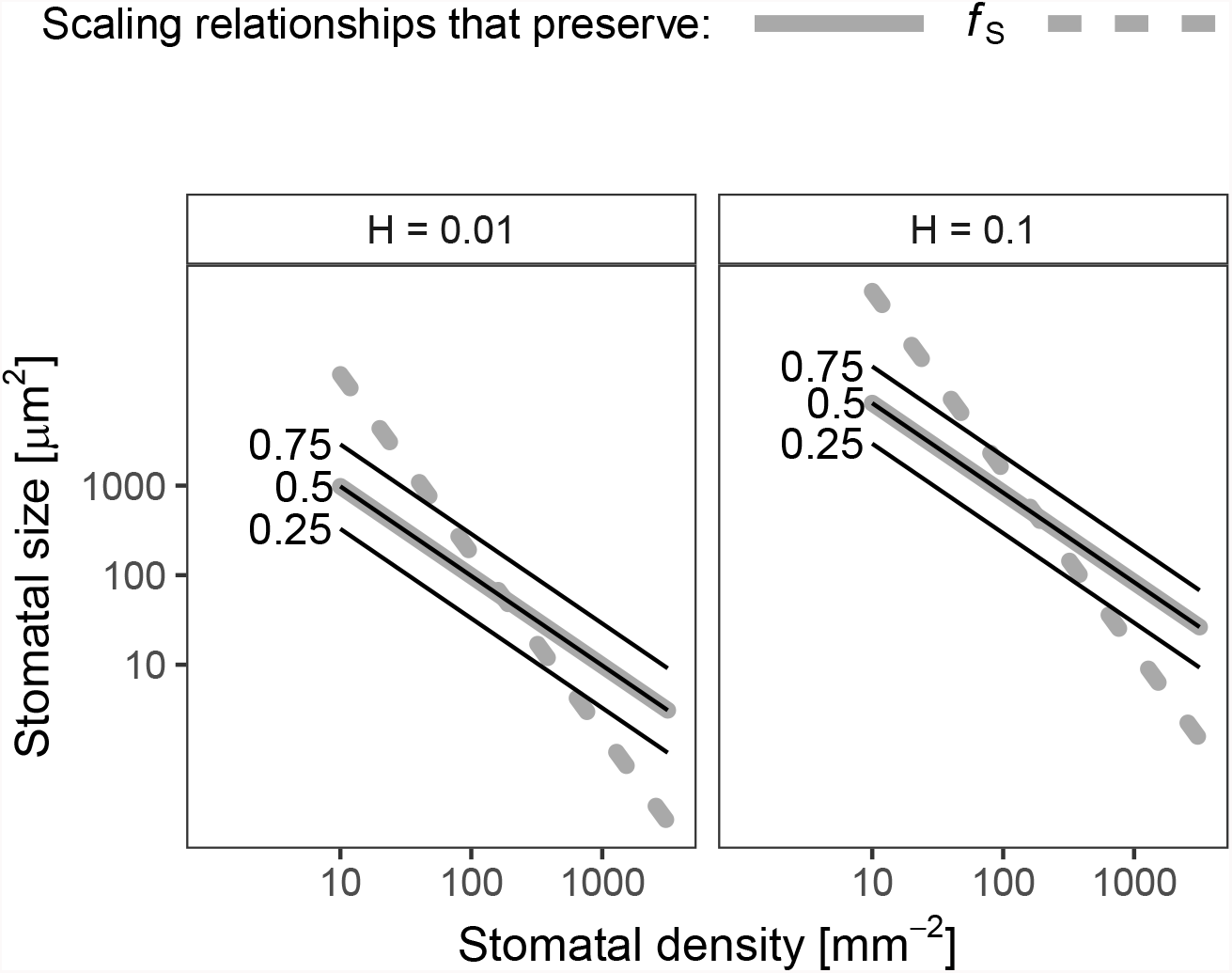
Log-log scaling relationships between stomatal density (*D*,*x*-axis) and size (*S*,*y*-axis) that preserve the probability of colonization (*p*_colonize_) in the spatially implicit model. As in Figure 4, in each panel, solid lines indicate values of *D* and *S* where *p*_colonize_ is 0.25 (lowest line), 0.5, or 0.75 (highest line). For reference, dashed grey lines show scaling relationships that preserve *f*_S_ (*β* = 1, slope = −1/*β* = −1) and *g*_s,max_ (*β* = 0.5, slope = −1/*β* = −2) drawn through the centroid of the plotting region. The scaling exponent is unity *β* = 1 when *H* > 0.

## APPENDIX 1 SPATIALLY IMPLICIT MODEL

A limitation of the spatially explicit model is that a pathogen could only infect stomata in the focal triangle where it landed. Here I analyze an alternative, spatially implicit, model that relaxes this assumptions. Instead, I assume that a pathogen can potentially infect any stomate on the leaf. It searches through a random walk and has a continuous, constant probability of encountering a stomate that is determined by stomatal cover (*f*_S_). If *fs* << 1, this can be modeled as a homogeneous Poisson process and the distance *x* a pathogen must travel before reaching a stomate follows an exponential distribution:

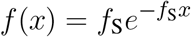

Given the constant death rate per unit distance *H*, the probability of surviving to distance *x* is *e*^−*Hx*^. The probability of locating a stomate the probability of surviving to distance *x* multiplied by *f*(*x*) and integrated over all *x* from 0 to *∞*:

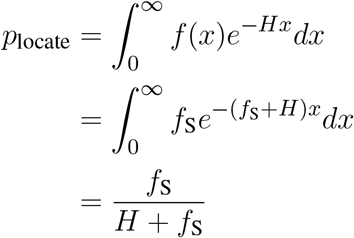

Substituting *p*_locate_ above into Equation 5:

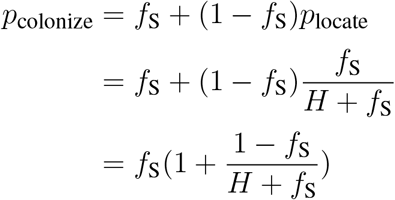

With the spatially implicit model, because pathogens can potentially reach any stomate on the leaf, *p*_colonize_ is greater than that in the spatially explicit model for the same value of *H*. For example, if the pathogen can search forever (*H* = 0), then it will always colonize (*p*_colonize_ = 1; Figure S2a). But even when *H* > 0, *p*_colonize_ is significantly higher than in the spatially implicit than spatially explicit model for the same *f*_S_ because pathogens can potentially colonize any stomate on the leaf.

Whereas the spatially explicit model probably underestimates *p*_locate_ for pathogens that can search over long distances, the implicit model overestimates because it assumes that the probability of encountering a stomate is constant (i.e. homogeneous Poisson process). This is not true because stomata are discrete areas on the leaf. If a pathogen is searching far away from a stomate, its probability of encountering a stomate in the near future is lower than that for a pathogen searching near a stomate. This should be modeled as a *nonhomogenous* Poisson process. Future work should derive *p*_locate_ for the stomatal anatomies presented here under a nonhomogeneous process.

Despite the quantitative differences in the the spatially explicit and implicit models, they have similar qualitative properties when *H* > 0, which is reasonable since the leaf surface is a relatively hostile environment for most pathogens (see Introduction). In both models, *p*_colonize_ increases with, but is higher than *f*_S_. In the spatially explicit model, size-density scaling that preserves *p*_colonize_ is 1 when *H* = 0 and slightly less than 1 otherwise (Figure 4). In the spatially implicit model, the scaling coefficient is always 1. Rearranging the equation for *p*_colonize_ above and substituting *f*_S_ = *DS*, the following relationship holds:

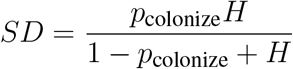

Since *H* is a constant the right-hand side of the equation above is constant for a given value of *p*_colonize_. Hence the *β_p_* that would preserve the relationship above is simply 1 (Figure S3).

