## Appendix 2 for "A stomatal model of anatomical tradeoffs between gas exchange and pathogen colonization"

### Appendix 2: Additional R packages

**Christopher D. Muir** [1*]

[1] *School of Life Sciences, University of Hawaii, Honolulu, Hawaii, USA*

Correspondence*:
Christopher D. Muir


| package | version | reference |
|---|---|---|
| assertthat | 0.2.1 | Wickham (2019a) |
| bibtex | 0.4.2.2 | Francois (2020) |
| BiocManager | 1.30.10 | Morgan (2019) |
| cli | 2.0.2 | Csárdi (2020) |
| codetools | 0.2-16 | Tierney (2018) |
| colorspace | 1.4-1 | Zeileis et al. (2019) |
| | | Zeileis et al. (2009) |
| | | Stauffer et al. (2009) |
| cowplot | 1.0.0 | Wilke (2019) |
| crayon | 1.3.4 | Csárdi (2017) |
| digest | 0.6.25 | Antoine Lucas et al. (2020) |
| dplyr | 1.0.0 | Wickham et al. (2020a) |
| ellipsis | 0.3.1 | Wickham (2020) |
| evaluate | 0.14 | Wickham and Xie (2019) |
| fansi | 0.4.1 | Gaslam (2020) |
| farver | 2.0.3 | Pedersen et al. (2020) |
| filehash | 2.4-2 | Peng (2006) |
| furrr | 0.1.0 | Vaughan and Dancho (2018) |
| future | 1.18.0 | Bengtsson (2020) |
| generics | 0.0.2 | Kuhn et al. (2018) |
| ggforce | 0.3.2 | Pedersen (2020) |
| ggimage | 0.2.8 | Yu (2020a) |
| gginnards | 0.0.3 | Aphalo (2019) |
| ggplot2 | 3.3.2 | Wickham (2016) |
| ggplotify | 0.0.5 | Yu (2020b) |
| globals | 0.12.5 | Bengtsson (2019a) |
| glue | 1.4.1 | Hester (2020) |
| gridGraphics | 0.5-0 | Murrell and Wen (2020) |
| gtable | 0.3.0 | Wickham and Pedersen (2019) |
| hms | 0.5.3 | Müller (2020) |
| htmltools | 0.5.0 | Cheng et al. (2020) |
| httr | 1.4.1 | Wickham (2019b) |
| jsonlite | 1.7.0 | Ooms (2014) |
| knitcitations | 1.0.10 | Boettiger (2019) |
| knitr | 1.29 | Xie (2020a) |
| | | Xie (2015) |

| package | version | reference |
|---|---|---|
| | | Xie (2014) |
| lifecycle | 0.2.0 | Henry (2020) |
| listenv | 0.8.0 | Bengtsson (2019b) |
| lubridate | 1.7.9 | Grolemund and Wickham (2011) |
| magick | 2.4.0 | Ooms (2020) |
| magrittr | 1.5 | Bache and Wickham (2014) |
| MASS | 7.3-51.6 | Venables and Ripley (2002) |
| munsell | 0.5.0 | Wickham (2018) |
| pillar | 1.4.6 | Müller and Wickham (2020a) |
| pkgconfig | 2.0.3 | Csárdi (2019) |
| plyr | 1.8.6 | Wickham (2011) |
| polyclip | 1.10-0 | Johnson and Baddeley (2019) |
| pracma | 2.2.9 | Borchers (2019) |
| purrr | 0.3.4 | Henry and Wickham (2020a) |
| R6 | 2.4.1 | Chang (2019) |
| Rcpp | 1.0.5 | Eddelbuettel and François (2011) |
| | | Eddelbuettel (2013) |
| | | Eddelbuettel and Balamuta (2017) |
| readr | 1.3.1 | Wickham et al. (2018) |
| RefManageR | 1.2.12 | McLean (2017) |
| | | McLean (2014) |
| rlang | 0.4.7 | Henry and Wickham (2020b) |
| rmarkdown | 2.3 | Allaire et al. (2020a) |
| | | Xie et al. (2018) |
| rticles | 0.14 | Allaire et al. (2020b) |
| rvcheck | 0.1.8 | Yu (2020c) |
| scales | 1.1.1 | Wickham and Seidel (2020) |
| sessioninfo | 1.1.1 | Csárdi et al. (2018) |
| stringi | 1.4.6 | Gagolewski (2020) |
| stringr | 1.4.0 | Wickham (2019c) |
| tibble | 3.0.3 | Müller and Wickham (2020b) |
| tidyr | 1.1.0 | Wickham and Henry (2020) |
| tidyselect | 1.1.0 | Henry and Wickham (2020c) |
| tikzDevice | 0.12.3.1 | Sharpsteen and Bracken (2020) |
| tweenr | 1.0.1 | Pedersen (2018) |
| units | 0.6-7 | Pebesma et al. (2016) |
| vctrs | 0.3.1 | Wickham et al. (2020b) |
| withr | 2.2.0 | Hester et al. (2020) |
| xfun | 0.15 | Xie (2020b) |
| xml2 | 1.3.2 | Wickham et al. (2020c) |
| yaml | 2.2.1 | Stephens et al. (2020) |
